# Regulation of Alternative Polyadenylation Events by PABPC1 Affects Erythroid Progenitor Cell Expansion

**DOI:** 10.1101/2025.03.17.643825

**Authors:** Yanan Li, Yanbo Yang, Bin Hu, Zi Wang, Wei Wang, Xiaofeng He, Xusheng Wu, Sheng Lin, Narla Mohandas, Hong Liu, Jing Gong, Long Liang, Jing Liu

## Abstract

Erythropoiesis is precisely regulated by multilayer networks and is crucial for maintaining steady-state hemoglobin levels and ensuring effective oxygen transport. Alternative polyadenylation (APA) is a post-transcriptional regulatory mechanism that generates multiple mRNA isoforms from a single gene, based on specific 3’-untranslated region sequences. While APA plays a vital role in various cellular processes, its mechanism in erythropoiesis remains unexplored. Here, we employed an integrative approach combining bioinformatics and experimental validation to systematically analyze APA’s role in erythropoiesis. We mapped the APA landscape during erythroid differentiation, discovering notable APA shifts crucial for the maturation from burst-forming unit erythroid to colony-forming unit erythroid. Notably, our findings highlighted *PABPC1* as the primary APA regulator of these stages. Functional investigations revealed that knocking down *PABPC1* disrupts erythroid progenitor cell proliferation and differentiation, implicating the protein’s essential role in modulating cell fate through APA regulation. We further identified that decreased *PABPC1* levels led to the increased usage of the proximal polyA site of *TSC22D1* and gene overexpression, revealing a novel mechanism through which APA affects erythroid progenitor expansion and differentiation. These insights uncover a novel dimension of APA regulation in early erythropoiesis, providing new strategies for the treatment of diseases associated with erythropoiesis disorders.

## Introduction

Hematopoietic stem cells (HSCs) generate megakaryocyte/erythrocyte progenitors through lineage commitment[1, 2]. These progenitors can differentiate into erythroid progenitors known as Burst-forming unit erythroid cells (BFU-Es) and colony-forming unit–erythroid (CFU-Es)[3, 4]. Then, erythroid progenitors undergo significant developmental changes, ultimately maturing into red blood cells[3, 5]. The expansion of erythroid progenitor cells plays a crucial role in limiting the efficacy of *in vitro* erythropoiesis induction[6]. The dysregulation of the proliferation is closely associated with various hematological disorders. For example, reduced expansion of erythroid progenitor cells has been observed in Diamond-Blackfan anemia (DBA) and is also recognized as a crucial factor contributing to the development of erythropoietin-unresponsive anemia[7]. Meanwhile, abnormal proliferation has been reported to be associated with diseases such as myelodysplastic syndromes (MDS), polycythemia vera (PV) and thalassemia[8–10]. Prior research has uncovered various molecules and pathways responsible for expansion of erythroid progenitor cells. For instance, glucocorticoids have been found to promote BFU-E expansion, whereas inhibiting *TGF-*β pathway and increased expression of RNA-binding protein *ZFP36L2* have both been demonstrated to enhance BFU-E proliferation[11, 12]. However, the specific mechanisms governing the expansion of erythroid progenitor cells and their regulation remain poorly understood[11].

Alternative cleavage and polyadenylation (APA), through the interactions of specific RNA-protein complexes and several key cis-elements such as polyadenylation signal (PAS) around the different polyA sites, could generate transcript isoforms with diverse 3’untranslated regions (3’UTRs)[13–15]. In previous studies, approximately 50-80% of mammalian pre-mRNA harbor multiple polyA sites[16, 17]. The widespread occurrence of APA not only enhances transcript complexity but also results in the gain or loss of binding sites for RNA-binding proteins and miRNAs, thereby influencing mRNA stability, localization, and translation[13, 14, 16, 18].

acvOver the past few years, research efforts have been directed toward understanding the regulation of APA in various biological processes, including neuronal differentiation, cell proliferation, and immune cell activation[13, 14, 16–23]. For example, in breast cancer, APA-mediated shortening of the 3’UTRs of *NRAS* and *c-JUN* was observed[24, 25]. Tumors with these shortened mRNA isoforms were also found to exhibit lower proliferation rates, yet display increased invasiveness[24, 25]. In cell proliferation, it has been found that increased cell activity can lead to widespread 3’UTR shortening[24, 25]. This process is a part of the gene expression program in cell proliferation, differentiation, and development[26]. APA core factors play a significant role in this process. In hematopoietic stem cells, for instance, *PABPN1*, a key regulator of APA, has been demonstrated to play a crucial role in regulating the genome-wide APA dynamics[27].

For erythropoiesis, research is less extensive, with only a few APA-related RBP studies reported[28, 29]. One such example is the APA core regulator *PABPC4*, has been identified to bind and stabilize transcript of *GPA*, along with other erythroid targets (*HBA1/HBA2, HBB, BTG2*, and *SLC4A1*)[28]. These interactions play a critical role in the maintenance of terminal erythroid maturation[28]. However, the global APA pattern during erythroid differentiation and their functions in the early stages of erythropoiesis remain largely unexplored.

In this study, we employed a combination of bioinformatics analysis and experiments to analyze global APA patterns across different stages of erythropoiesis. This approach enabled us to precisely capture and map the dynamic APA landscape during erythroid differentiation, pinpointing key APA shifts pivotal for the transition from BFU-E to CFU-E stages. Our findings revealed a marked downregulation of the APA factor *PABPC1* during these stages, which might precipitate notable APA alterations. Furthermore, our investigations identified *TSC22D1* as a crucial target influenced by *PABPC1*-mediated APA events. Changes in *PABPC1* levels were observed to promote the usage of its proximal polyA site and lead to an increase in *TSC22D1* expression. In-depth analysis demonstrated that *TSC22D1* plays a crucial role in the expansion and differentiation of erythroid progenitor cells. In summary, this discovery provides a clearer understanding of the molecular mechanisms of APA at early stages of erythropoiesis and underscores potential therapeutic targets for disorders related to abnormal erythroid cell development.

## Results

### Global APA Patterns in Erythropoiesis

Efficient regulation of erythropoiesis is crucial for maintaining systemic homeostasis[7–10]. To explore APA’s influence on erythroid differentiation, we retrieved RNA-Seq data from our previous research, which contains both early (GSE61566) and late (GSE53983) stages of erythropoiesis, spanning seven phases from BFU-E to orthochromatic erythroblasts (Figure 1A). We further employed DaPars v2.0 to systematically estimate the relative usage of APA on these datasets[30]. DaPars v2.0 employs a two-normal mixture model, enabling quantification of APA usage across multiple samples (https://academic.oup.com/nar/article /50/D1/D39/6357732) (Figure 1B). In total, we identified 8,377 APA events during erythropoiesis. PAS motifs are the key cis-elements for polyadenylation and play crucial roles in the processing and stabilization of mRNA[31, 32]. To investigate the preferred PAS motifs of these APA events, we extracted sequences of ±50 nt around each proximal and distal polyA sites and employed DREME to enrich PAS features. Our study revealed a preference for the usage of AATAAA and ATTAAA at the distal polyA sites, which are known as canonical PAS motifs, while the proximal polyA sites showed a tendency towards AATAAA and AAGAAA. AAGAAA is also recognized as an active polyadenylation signal in mammalian cells[33] (Figure 1C). To validate biological replicability of PDUI values, we conducted Pearson correlation analyses on the PDUI matrix of erythropoiesis. We observed clear stage-specific clustering, indicating a significant variation in polyA site utilization across different erythropoiesis stages (Figure 1D).

**Figure 1.**
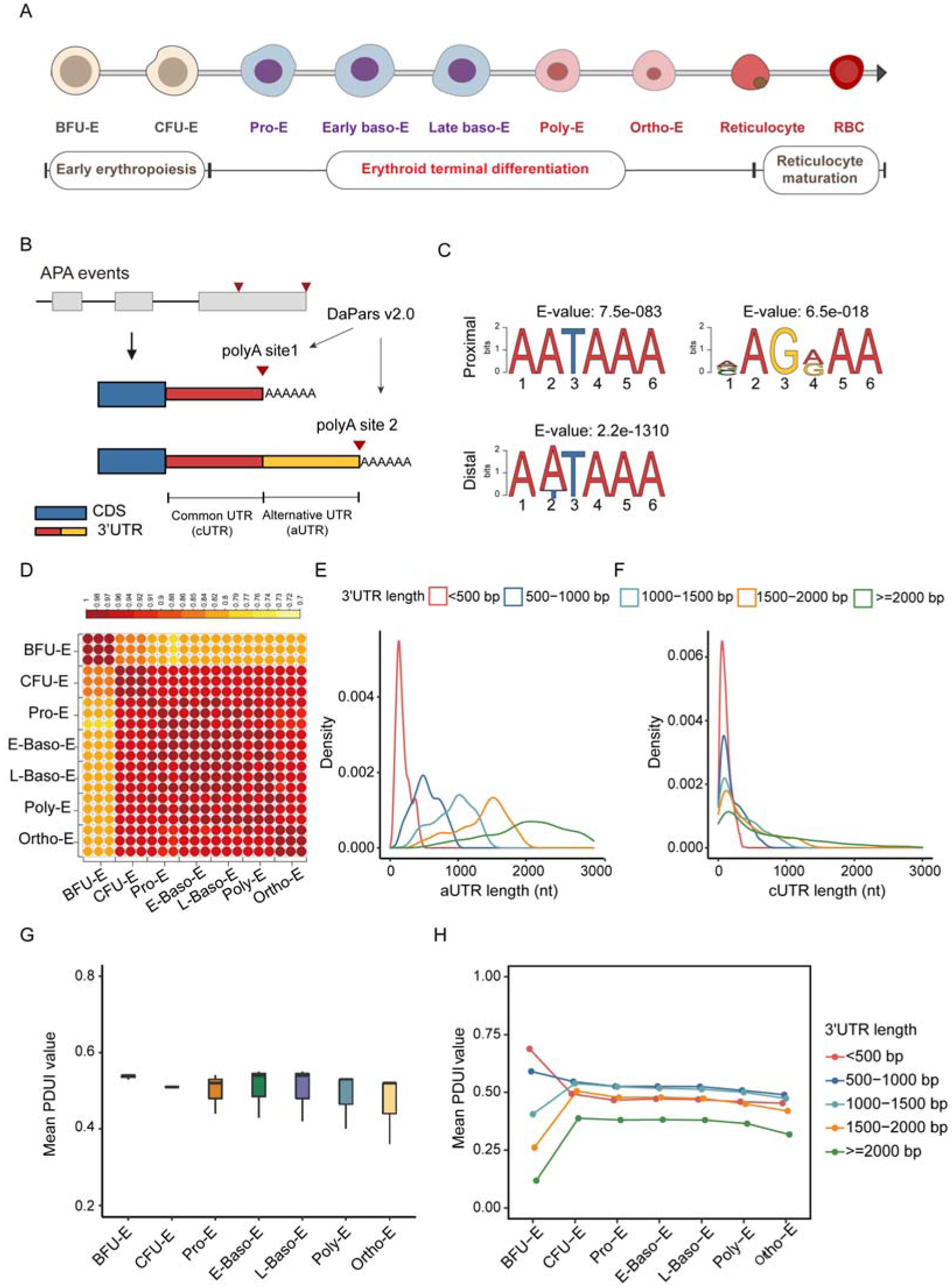
Global Patterns of Alternative Polyadenylation Events in Erythropoiesis. **A.** Schematic diagram of different stages of human erythroid differentiation. The development process of erythropoiesis includes three stages early erythropoiesis, terminal erythroid differentiation and reticulocyte maturation. **B**. Schematic of APA isoforms using proximal PAS (pPAS) or distal PAS (dPAS) in the 3’UTR. The region between the PASs is named alternative 3’UTR (aUTR). The common region of isoforms is named common 3’UTR (cUTR). **C**. Enriched motifs around proximal and distal polyA sites of APA events. **D**. Pearson correlation heatmap displaying the PDUI across erythropoiesis stages. The color of each cell represents the correlation coefficient, with dark red signifying a strong positive correlation and yellow denoting a weaker correlation. **E**. Density distribution of alternative UTR lengths across various 3’UTR length categories. The color indicates the stratification of 3’UTR lengths into different categories. **F**. Density distribution of common UTR lengths across various 3’UTR length categories. The color indicates the stratification of 3’UTR lengths into different categories. **G**. The average relative 3’UTR length at each stage of human erythropoiesis (BFU-E to Ortho-E). **H**. Mean PDUI values across different erythroid maturation stages for 3’UTR length categories. The x-axis indicates the erythroid maturation stages from BFU-E to Ortho-E, while the y-axis shows the mean PDUI values. Each line represents a distinct 3’UTR length category.

We further investigated the structural characteristics of APA events during erythropoiesis, and found that the 3’UTR lengths of APA events were significantly longer than those of other genes (Wilcoxon test, *P* < 2.2e−16) (Figure S1A). To explore the finer structural differences in different 3’UTR lengths, we classified the 3’UTR lengths into distinct bins (<500bp, 500-1000bp, 1000-1500bp, 1500-2000bp, ≥2000bp), and discovered that APA events were predominantly located in the shorter 3’UTR length bins, particularly within the <500 bp category (Figure 1E and 1F). aUTR (alternative UTR) is defined as the portion of the 3’UTR that exhibits length variations, and cUTR (common UTR) remains consistent across different isoforms[34]. We also observed that longer 3’UTRs correlate with longer aUTRs (Figure 1E and S1B), while the cUTR remains relatively stable across different bin types (Figure 1F and S1C). As 3’UTRs contain a variety of cis-elements, such as miRNA target sites and RBP binding sites. Longer 3’UTRs and aUTRs imply an increased probability of variability in these regulatory elements, thereby influencing gene expression and cellular functions.

Next, we investigated APA dynamics during erythropoiesis, and noticed a decrease in PDUI in the early phases of erythropoiesis (Figure 1G), suggesting an enhanced utilization of proximal polyA sites from BFU-E to CFU-E stages. Additionally, APA events with shorter 3’UTRs (< 1000 bp) exhibit an increased utilization of proximal polyA sites, whereas longer 3’UTRs are more likely to undergo a transition toward distal polyA site selection (Figure 1H). These findings suggest a complex regulatory role of APA and highlight its potential biological significance in erythroid early progenitor cells.

### 3’UTR shifting of genes during Erythropoiesis

To investigate genes that undergone significant 3’UTR changes during erythropoiesis, we compared PDUI values between different stages, and identified 5,435 genes exhibit 3’UTR lengthening or shortening. We noticed a pronounced trend where APA events with 3’UTR shortening considerably outnumber those with 3’UTR lengthening during erythropoiesis. These APA alterations, which included both 3’UTR lengthening and shortening, primarily occur within the BFU-E to Pro-E stages and tend to stabilize in the late stages of erythropoiesis (Figure 2A). To elucidate the functional roles of these genes, we categorized them based on the 3’UTR lengthening or shortening, and subsequently conducted the gene enrichment analysis. These APA events at different stages are mainly enriched in pathways associated with cell cycle and RNA regulation. Specifically, during the transition from BFU-E to CFU-E stages, pathways such as ‘Cell cycle’ and ‘Metabolism of RNA’ are the most significantly enriched among the shortening APA events. Conversely, during the transition from CFU-E to Pro-E stages, the lengthening APA events show the highest enrichment in pathways such as ‘Mitotic cell cycle’ and ‘Protein localization to organelle’, indicating a dynamic interplay between APA event regulation and erythropoiesis (Figure 2B).

**Figure 2.**
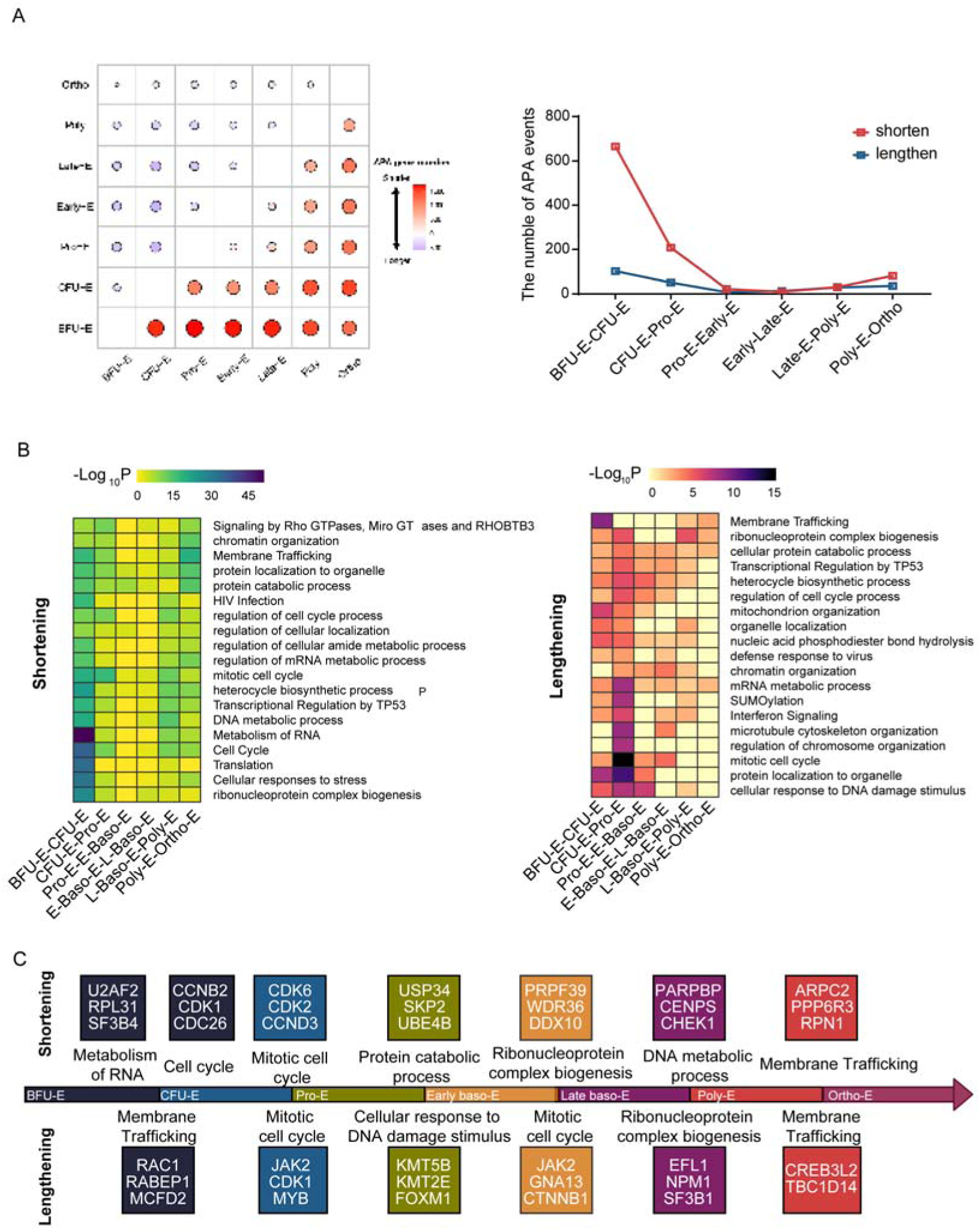
3’UTR shifting of genes during Erythropoiesis. A. The number of shortening or lengthening APA events in each stage of human erythropoiesis (BFU-E to Ortho-E). Left graph shows the number of genes at different erythroid differentiation stages (from BFU-E to Ortho-E) that exhibit changes in 3’UTR length, including lengthening and shortening. Right graph displays the variation in the number of APA events from BFU-E to Poly-E stages, where the red line indicates the number of genes with shortened 3’UTRs and the blue line shows those with lengthened 3’UTRs. **B**. Heatmap of gene enrichment results of APA patterns (shortening APA events and lengthening APA events) of erythropoiesis. Left graph lists biological processes associated with 3’UTR shortening. Right graph lists biological processes associated with 3’UTR lengthening. **C**. Example diagram of enriched pathway related genes in various stages of erythropoiesis. Upper half lists key genes associated with 3’UTR shortening and related biological processes. Lower half lists key genes associated with 3’UTR lengthening and related biological processes.

We further focused on the representative APA genes which are from the specific gene sets and have been proved essential to erythropoiesis. During the BFU-E to CFU-E stages, the representative genes were observed in pathways such as RNA metabolism (*U2AF2, RPL31,* and *SF3B4*), the cell cycle (*CCNB2, CDK2,* and *CDC26*), and membrane transport (*RAC1, RABEP1,* and *MCFD2*). In the terminal stages of erythropoiesis, these genes are enriched in mitotic cell cycle (*CDK6, CDK2, CCND3, JAK2, CDK1,* and *MYB*), biogenesis of ribonucleoprotein complexes (*PRPF39, WDR36, DDX10, EFL1, NPM1,* and *SF3B1*), membrane transport (*ARPC2, PPP6R3, RPN1, CREB3L2,* and *TBC1D14*), and protein metabolic processes (*USP34, SKP2,* and *UBE4B*) (Figure 2C). These findings suggest that APA dynamics are functionally diverse across various erythropoiesis phases, with a marked association with cell cycle and RNA regulation, particularly in early erythropoiesis.

### PABPC1 as a Potential APA Regulator for 3’UTR Shortening during BFU-E to CFU-E Stages

RNA-binding proteins are crucial in the regulation of APA by binding to specific regions within the 3’UTR, consequently influencing the PAS choice[13, 16]. Prior studies have proved the significant roles of RBPs in cellular development, differentiation and cancers[13, 14, 16, 18–23]. To investigate the principal factors regulating genes exhibiting 3’UTR lengthening or shortening during the early stages of erythropoiesis, we first collected a series of known core APA factors. Subsequently, we calculated the correlation between PDUI of each APA event and expression of each APA factor. Significant associations between known 3’ end processing factors and APA events were obtained using stringent criteria (R^2^≥ 0.3 and FDR < 0.05). After aggregating the association results by RBP, we found the top 10 factors potentially mediating APA events were *PABPC1, PAPOLG, CPSF2, CSTF2T, FIP1L1, WDR33, CPSF7, PPP1CA, PABPN1* and *CPSF6* (Figure 3A). To ascertain whether these factors exhibit genuine variability throughout the differentiation process, thereby elucidating their potential roles in erythropoiesis, we calculated the standard deviation of the expression levels (measured in TPM) of these factors across all samples of erythrocyte differentiation. Our findings demonstrated that *PABPC1* exhibited the highest standard deviation and was markedly higher than that of other factors. This observation indicates a considerable fluctuation in *PABPC1* expression throughout erythropoiesis, highlighting its potential involvement in the regulatory mechanisms that govern the erythropoiesis (Figure 3B). Considering the significant APA changes during the early stages of erythropoiesis, we quantified the mRNA levels of these genes during these critical stages. Remarkably, only *PABPC1* exhibited relatively high expression in the stage of BFU-E (Figure 3C). Furthermore, we induced CD34+ cells derived from human mobilized peripheral blood samples to differentiate into the erythroid lineage[3]. We found that both mRNA and protein expression levels of *PABPC1* were the highest during the BFU-E stage and saw a decline in the CFU-E stage (Figure 3D and E). Among the APA events associated with *PABPC1*, we observed 1,276 events leading to 3’UTR shortening and 243 events resulting in 3’UTR lengthening, indicating knockdown of *PABPC1* may facilitate an increased usage of proximal polyA sites. PABPC1 has been shown to play a role in promoting distal site usage, with knockout shortening the 3’UTR[35], however, such observations have not been reported in the erythroid lineage. These results suggest that *PABPC1* may be a crucial APA regulator mediating the 3’UTR shortening of transcripts from the BFU-E to CFU-E stages during erythropoiesis.

**Figure 3.**
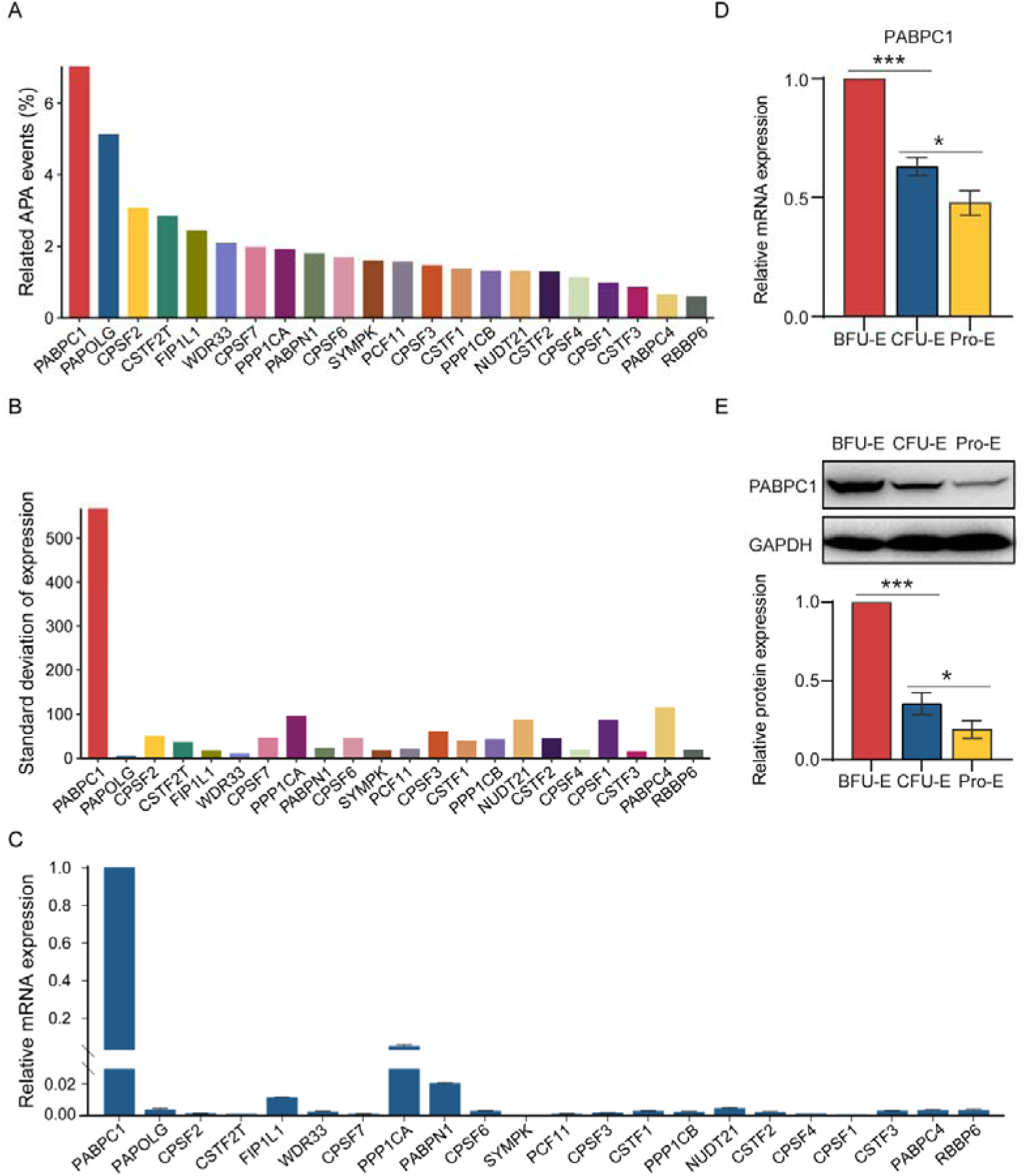
PABPC1 as a Potential APA Regulator for 3’UTR Shortening during BFU-E to CFU-E Stages. A. A bar chart showing the ability of APA regulatory factors to regulate APA events during erythroid development. The vertical axis represents the proportion of APA-RBP associations relative to all APA events with 3’UTR lengthening or shortening during differentiation, which reflects the ability of core factors to regulate APA events. The horizontal axis represents the core APA regulatory factors. **B**. The standard deviation of the expression levels of core APA factors across all samples of erythrocyte differentiation. **C**. RT-qPCR shows the mRNA expression of core APA factors relative to PABPC1 in early erythropoiesis (BFU-E stage). Cells from the BFU-E stage were sorted after human mobilized peripheral blood CD34+ cells were induced to differentiate into the early erythroid lineage in vitro. **D**. RT-qPCR shows the mRNA expression of PABPC1 from BFU-E to Pro-E stage. Cells at stages from BFU-E to Pro-E were sorted following in vitro differentiation of human mobilized peripheral blood CD34+ cells into the erythroid lineage. **E**. Representative images of western blotting showing PABPC1 expression from BFU-E to Pro-E stage. Cells at stage from BFU-E to Pro-E were sorted following in vitro differentiation of human mobilized peripheral blood CD34+ cells into the specific erythroid lineage. Quantitative analysis of PABPC1 protein expression data from three independent experiments from BFU-E to Pro-E stage. GAPDH was used as a loading control.

### PABPC1 Plays a Crucial Role in the Development from the BFU-E to CFU-E Stage, Impacting Proliferation and Differentiation

*PABPC1* is widely expressed in most eukaryotes[36] This factor interacts with various proteins and exhibits complex intracellular localization pattern[37]. In mammals, *PABPC1* has been demonstrated to be a multifunctional RNA-binding protein, indispensable for protein translation initiation, RNA processing, RNA stability, and the promotion of distal polyA utilization[36, 38]. Under normal conditions, *PABPC1* is predominantly localized in the cytoplasm[37, 39]. However, under specific circumstances, it has been proved to shuttle between the nucleus and cytoplasm, where it polyadenylates intron-containing precursor mRNAs and interacts with polyA polymerase, participating in nuclear RNA processing[37, 39].

To further investigate the role of *PABPC1* in early human erythropoiesis and analyze its functions, we employed an shRNA lentivirus to knockdown *PABPC1* expression in the initial stages of erythropoiesis in vitro (Figure 4A, B, and C). We conducted cell counting and generated growth curves to evaluate the effect of *PABPC1* knockdown on cell proliferation during early erythropoiesis, revealing a significant inhibition in cell proliferation following *PABPC1* knockdown (Figure 4D). Next, we employed flow cytometry to examine the effect of *PABPC1* knockdown on early human erythropoiesis. The results for early erythropoiesis indicated that the proportion of CD34− CD36+ CD71+ (CFU-E) cells markedly decreased, while the proportion of CD34+ CD36− CD71− (BFU-E) cells remained largely unchanged after *PABPC1* knockdown (Figure 4E and F). Similarly, the colony formation experiment revealed a significant reduction in the number of CFU-E colonies post-*PABPC1* knockdown, while the number of BFU-E colonies showed no significant change (Figure 4G). Additionally, we evaluated cell cycle and apoptosis using flow cytometry (Figure 4H-K). The outcomes revealed a significant increase in cell apoptosis rate and cell cycle arrest in the G0/G1 phase after *PABPC1* knockdown (Figure 4H-K). Moreover, we constructed overexpression vectors for *PABPC1* to investigate its functional role. To further explore the contribution of *PABPC1* in early human erythropoiesis, we overexpressed it during the initial stages of erythroid differentiation in vitro using CD34^+^ hematopoietic stem cells. Flow cytometry analysis revealed a significant increase in the proportion of CD34^−^ CD36^+^ CD71^+^ cells (CFU-E), while the population of CD34^+^ CD36^−^ CD71^−^ cells (BFU-E) remained largely unchanged following *PABPC1* overexpression (Figure S2A and S2B). Additionally, colony formation assays demonstrated a notable increase in the number of CFU-E colonies upon *PABPC1* overexpression, whereas BFU-E colony formation showed no significant alteration (Figure S2C). Given the reported importance of *PABPN1* and *PABPC4*, we also evaluated their mRNA expression levels. Our results indicated that both *PABPN1* and *PABPC4* were expressed at relatively low levels (Figure S3A). Notably, *PABPN1* expression remained stable during early erythroid development (Figure S3A), while *PABPC4* mRNA levels exhibited a significant increase from the BFU-E to CFU-E stage despite its overall low expression (Figure S3A). Based on this finding, we focused on *PABPC4* for functional studies and observed that its knockdown did not significantly affect erythroid progenitor proliferation or differentiation (Figure S3C-E). Taken together, these findings highlight the critical role of *PABPC1* in regulating the expansion and transition of erythroid progenitor cells.

**Figure 4.**
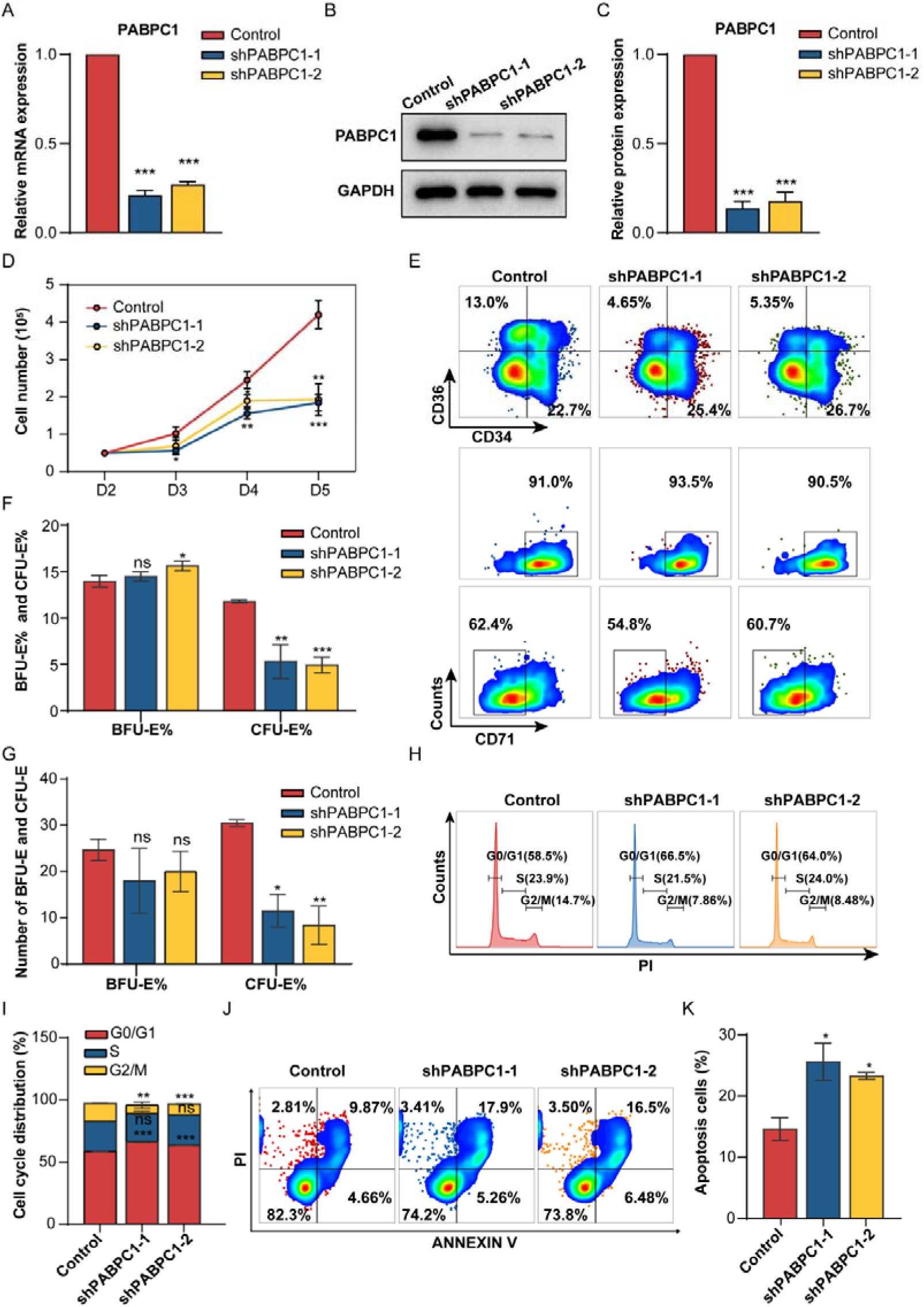
PABPC1 Plays a Crucial Role in the Development from the BFU-E to CFU-E Stage, Impacting Proliferation and Differentiation. A. RT-qPCR shows PABPC1 mRNA expression in early erythroblasts infected with lentivirus containing control shRNA or PABPC1 shRNA. **B**. Representative images of western blotting analysis showing PABPC1 expression in erythroblasts infected with lentivirus containing control shRNA or PABPC1 shRNA. **C**. Quantitative analysis of protein expression data from three independent experiments of (B). GAPDH was used as a loading control. **D**. Early erythroid cell (BFU-E) growth curves determined by cell counting. early erythroid cell following infection with the same amount of lentivirus as indicated in (A). Statistical analysis of the data from three independent experiments; the bar plot represents the mean ± SD of triplicate samples. *P<0.05, **P<0.01, ***P<0.001 versus control based on Student’s t-test. **E**. Representative flow cytometry data of early differentiation for early erythroid cell (BFU-E) infected with the same amount of lentivirus as indicated in (A). CD34- CD36+ CD71high represent the stage of CFU-E cells, while CD34+ CD36- CD71low represent the stage of BFU-E cells. CD34- CD36+ CD71high and CD34+ CD36- CD71low cells gate from CD45+ GPA- CD123- cells. **F**. Statistical analysis of the CD34- CD36+ CD71high (CFU-E) and CD34+ CD36- CD71low (BFU-E) cells rate (%) from three independent experiments is shown. **G**. Statistical analysis of the number of BFU-E and CFU-E formed in soft agar clone formation experiments. **H**. Representative flow cytometry data of cell cycle for early erythroid cell (BFU-E) infected with the same amount of lentivirus as indicated in (A). **I**. Statistical analysis of the PI fluorescence intensity (represent each stage of cell cycle, G0/G1, S, G2/M respectively) from three independent experiments is shown. **J**. Representative flow cytometry data of cell apoptosis for early erythroid cell (BFU-E) infected with the same amount of lentivirus as indicated in (A). **K**. Statistical analysis of the apoptosis cells rate (%) from three independent experiments is shown.

### PABPC1 Knockdown Significantly Affects APA Events from BFU-E to CFU-E During Early Erythropoiesis

To explore *PABPC1*’s transcriptional role in early erythropoiesis, we conducted *PABPC1* knocked out at the early stage of erythropoiesis and performed nanopore sequencing (Figure 5A). Nanopore third-generation full-length transcriptome can obtain full-length transcripts without the need for interruption and splicing, accurately identifying transcript structural information such as alternative splicing (AS), fusion genes, and non-coding RNA. At the same time, it can directly perform quantitative analysis at the gene/transcript level. The ONT direct RNA sequencing method can also detect multiple base modifications without amplification, reducing the base preference introduced by the PCR process[40, 41].

**Figure 5.**
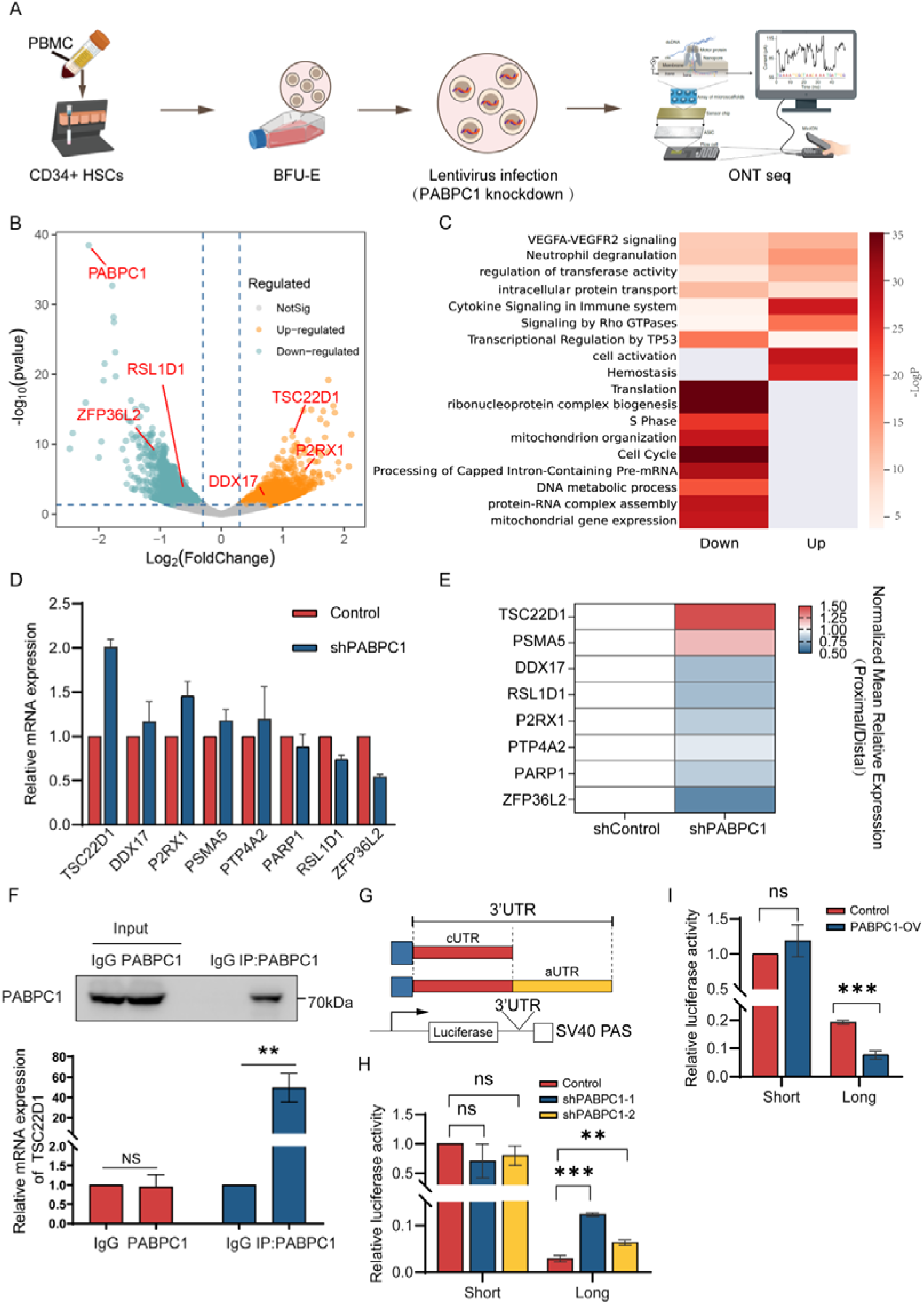
PABPC1 Knockdown Significantly Affects APA Events from BFU-E to CFU-E During Early Erythropoiesis. A. Flow chart of sample treatment and collection for nanopore sequencing. **B**. Differential gene analysis of nanopore sequencing data from early erythrocyte which knocking down PABPC1. Blue circles represent downregulated genes and yellow circles represent upregulated genes. **C**. Gene Ontology (GO) enrichment analysis on differential expression genes (PABPC1 knockdown vs control) in early erythropoiesis. **D**. The validation of mRNA expression of differential expression genes by RT-qPCR. **E**. Heatmap representing normalized mean relative expression of the proximal to the distal mRNA isoform based on qRT-PCR analysis (PABPC1 knockdown vs control). Results > 1 indicate relative increase of the proximal transcript, whereas results < 1 indicate relative increase of the distal transcript. **F**. Validation of antibody validity by IP-Western blot assay. Lane 1 and Lane 2: Input; Lane 3, IgG negative control; Lane 4, IP sample using anti-PABPC1 antibody (above panel). A RIP assay was performed using anti-normal rabbit IgG or anti-PABPC1 in BFU-E cell lysates. The relative expression level of TSC22D1 was detected by qRT-PCR (below panel). **G**. Schematic of APA isoforms using proximal PAS (pPAS) or distal PAS (dPAS) in the 3’UTR. The region between the PASs is named alternative 3’UTR (aUTR). The common region of isoforms is named common 3’UTR (cUTR). The proximal or distal regulatory sequences of the TSC22D1 gene 3’ UTR were cloned into the PGL3-BASIC reporter vector and validated by sequencing. **H**. Luciferase reporter assays to evaluate the impact of PABPC1 knockdown on the regulatory efficiency of functional variants in the proximal or distal regions of the TSC22D1 gene 3’ UTR. Luciferase activity was measured using a dual-luciferase detection system. Quantitative analysis was conducted by calculating the Fluc/Rluc ratio. **I**. Luciferase reporter assays to evaluate the impact of PABPC1 overexpression on the regulatory efficiency of functional variants in the proximal or distal regions of the TSC22D1 gene 3’ UTR. Luciferase activity was measured using a dual-luciferase detection system. Quantitative analysis was conducted by calculating the Fluc/Rluc ratio.

We employed LAPA to perform a comprehensive global analysis of alternative polyadenylation (APA) using ONT data (see Methods). Our analysis revealed that *PABPC1* knockdown consistently increased proximal polyadenylation site usage across different thresholds (Table S1), suggesting that *PABPC1* deficiency shifts polyadenylation toward proximal sites. For each APA-associated gene, we compared the median polyA tail lengths at proximal and distal sites (File S1). Across all ONT samples, distal isoforms exhibited longer polyA tails (Figure S4A and Table S2), with an overall median length of 27 nucleotides compared to 24 nucleotides at proximal sites. Additionally, *PABPC1* knockout was associated with polyA tail shortening (Figure S4B), underscoring its role in tail length maintenance. Notably, in *PABPC1*-depleted cells, polyA tail shortening was primarily linked to increased proximal site usage (Figure S4C). This suggests that *PABPC1* may simultaneously regulate both polyA tail length and APA site selection, with its absence tipping the balance toward proximal site usage and shorter polyA tails.

On the other hand, our differential gene expression analysis identified 1,696 upregulated and 2,042 downregulated genes, with *PABPC1* itself markedly downregulated (log2FoldChange=-2.16, P-value=3.45e-39) (Figure 5B). Subsequently, the gene ontology analysis revealed divergent regulatory trends. The upregulated genes were predominantly linked to cell activation, hemostasis, and cytokine signaling in the immune system, whereas the downregulated genes were associated with the cell cycle, translation, and ribonucleoprotein complex biogenesis (Figure 5C).

Due to the crucial role of the pathways such as cell cycle and cellular activation in both APA and erythropoiesis, we selected a set of genes differentially expressed in these biological processes to examine their expression and APA changes. Under *PABPC1* knockdown conditions, our qRT-PCR analysis indicated upregulation in *TSC22D1, DDX17,* and *P2RX1*, and downregulation in *ZFP36L2* and *RSL1D1*, consistent with our RNA-seq results (Figure 5D).

For APA analysis, we quantified the proximal and distal polyA sites using qRT-PCR and calculated the relative usage of proximal polyA sites. We observed a general increase in the use of distal polyA sites was noted for genes such as *DDX17, P2RX1*, and *ZFP36L2*. In contrast, *TSC22D1* and *PSMA5* exhibited a significant increase in proximal polyA site usage, which corresponded with increase expression (Figure 5E). The IGV snapshots show increased proximal polyA site usage in *TSC22D1* and APA alterations in other significant genes (Figure S5A-D). To validate the detectable signal at the distal site, we quantified polyA site usage through qRT-PCR. *PABPC1* knockdown (KD) resulted in a significant increase in proximal polyA site usage with a corresponding decrease in distal site usage (Figure S5E). To comprehensively assess *PABPC1’s* regulatory role in APA, we employed both knockdown and overexpression strategies. RIP assays confirmed *TSC22D1* mRNA-*PABPC1* protein interaction in erythroid progenitor cells, with significantly higher mRNA enrichment in the *PABPC1* group versus the IgG control (Figure 5F). We constructed firefly luciferase reporters containing either: 1) *TSC22D1* constitutive 3′UTRs (cUTRs, corresponding to proximal PAS usage), or 2) extended 3′UTRs incorporating both cUTR and alternative regions (aUTRs) from the longer APA isoform (Figure 5G). To standardize transcript processing, all constructs were terminated with a heterologous SV40 PAS (Figure 5G). Following transfection of equimolar reporter constructs into erythroid progenitor cells, we measured luciferase activity to evaluate APA-mediated regulatory effects. Notably, luciferase expression driven by proximal PAS-derived 3′UTRs remained stable upon *PABPC1* manipulation (Figure 5H-I and S6A-B). In contrast, the extended 3′UTR displayed significantly reduced luciferase activity with *PABPC1* overexpression, whereas *PABPC1* KD enhanced its activity (Figure 5H-I and S6A-B). These findings demonstrate that *PABPC1* is essential for maintaining distal polyA site selection in *TSC22D1*, thereby modulating *TSC22D1* expression through this mechanism.

### PABPC1 Mediates Expansion and the Transition from BFU-E to CFU-E by Influencing *TSC22D1* APA Changes

The *TSC22D* protein family encompasses a range of activities involved in the regulation of cell proliferation and differentiation[42, 43]. Among these family members, *TSC22D1*, the first identified member, is well-known for its tumor suppressor function[44]. For instance, in cervical cancer cells, knockdown of the RNA-binding protein *MEX3D* has been shown to inhibit cell proliferation and colony formation while promoting apoptosis by stabilizing *TSC22D1* mRNA[44].

To investigate whether PABPC1 influences the proliferation of early erythroid progenitor cells by modulating *TSC22D1*, we examined *TSC22D1* expression from the BFU-E to Pro-E stages. Both database analysis and qPCR verification indicated a significant increase in *TSC22D1* mRNA levels from the BFU-E to CFU-E stages, a trend opposite to that of *PABPC1* (Figure 3D, 6A and S7A). Western blot assays corroborated this pattern at the protein level (Figure 3E 6B and 6C), suggesting a negative regulation of *TSC22D1* expression by *PABPC1*.

Subsequently, we performed *PABPC1* knockdown followed by *TSC22D1* knockdown. *PABPC1* knockdown led to an increase in *TSC22D1* expression, but subsequent *TSC22D1* knockdown counteracted this increase without affecting *PABPC1* expression (Figure 6D, E and F). Further investigations revealed that *PABPC1* knockdown increased *TSC22D1* expression, significantly reduced cell proliferation, hindered differentiation (notably decreasing CFU-E cells), increased cell apoptosis, and induced cell cycle arrest at the G0/G1 phase. These changes could be rescued by *TSC22D1* knockdown, which restored *TSC22D1* levels after *PAPBC1* knockdown (Figures 6G-L). Therefore, these results indicate that *PABPC1* changes the ability of cells expansion and differentiation by affecting the APA alteration of *TSC22D1*.

**Figure 6.**
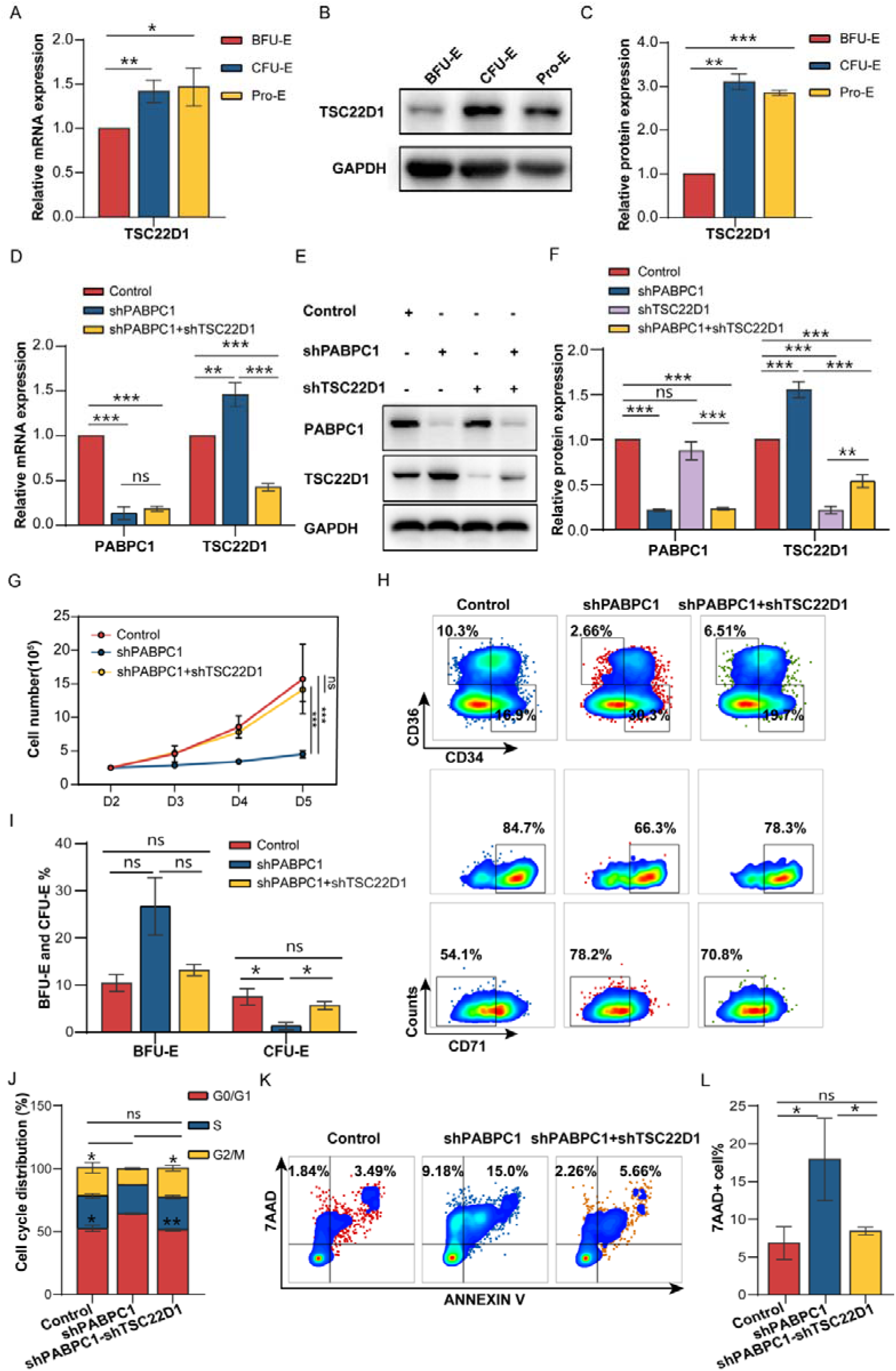
PABPC1 Mediates Expansion and the Transition from BFU-E to CFU-E by Influencing *TSC22D1* APA Changes. A. RT-qPCR shows the mRNA expression of TSC22D1 in early erythropoiesis. **B**. Representative images of western blotting analysis showing TSC22D1 expression in early erythropoiesis. **C**. Quantitative analysis of TSC22D1 protein expression data from three independent experiments in early erythropoiesis. GAPDH was used as a loading control. **D**. RT-qPCR shows PABPC1 and TSC22D1 mRNA expression in early erythroblasts infected with lentivirus containing control shRNA, PABPC1 shRNA or PABPC1 shRNA+ TSC22D1 shRNA. **E**. Western blotting shows PABPC1 and TSC22D1 protein expression in early erythroblasts infected with lentivirus containing control shRNA, PABPC1 shRNA or PABPC1 shRNA+ TSC22D1 shRNA. **F**. Quantitative analysis of PABPC1 and TSC22D1 protein expression data from three independent experiments in early erythropoiesis. GAPDH was used as a loading control. **G**. Early erythroid cell (BFU-E) growth curves determined by cell counting early erythroid cell following infection with the same amount of lentivirus as indicated in (D). Statistical analysis of the data from three independent experiments; the bar plot represents the mean ± SD of triplicate samples. *P<0.05, **P<0.01, ***P<0.001 versus control based on Student’s t-test. **H**. Representative flow cytometry data of early differentiation for early erythroid cell (BFU-E) infected with the same amount of lentivirus as indicated in (E). CD34- CD36+ CD71high represent the stage of CFU-E cells, while CD34+ CD36- CD71low represent the stage of BFU-E cells. CD34- CD36+ CD71high and CD34+ CD36- CD71low cells gate from CD45+ GPA- CD123- cells. **I**. Statistical analysis of the CD34- CD36+ CD71high (CFU-E) and CD34+ CD36- CD71low (BFU-E) cells rate (%) from three independent experiments is shown. **J**. Statistical analysis of the PI fluorescence intensity (represent each stage of cell cycle, G0/G1, S, G2/M respectively) from three independent experiments is shown. **K**. Representative flow cytometry data of cell apoptosis for early erythroid cell (BFU-E) infected with the same amount of lentivirus as indicated in (E). **L**. Statistical analysis of the apoptosis cells rate (%) from three independent experiments is shown.

## Discussion

Erythropoiesis is tightly regulated to maintain homeostasis. Any disruption in this process can lead to blood disorders such as polycythemia vera, thalassemia, and myelodysplastic syndromes[7–10]. However, the understanding of the mechanisms underlying the erythroid lineage differentiation, especially during the early stages, are remains limited.

Previous studies have documented APA’s critical role in cell differentiation, particularly affecting pluripotent stem cells and early progenitor cell proliferation[28, 29]. Although APA’s importance in the erythroid system has been reported[27], a systematic analysis of APA’s dynamic changes within this system is still lacking.

Our study focuses on the role of APA in erythropoiesis. We applied the DaPars v2.0 algorithm to RNA-Seq data from different erythroid developmental stages. We discovered significant APA changes during the transition from BFU-E to CFU-E, which are early stages in erythroid progenitor development. This finding indicates that APA is crucial for the growth and self-renewal of erythroid progenitor cells. In addition, by analyzing the proportion of core APA regulatory factors and combining this with gene expression experiment, we identified polyadenylate-binding protein cytoplasmic 1 (*PABPC1*) as a crucial regulator of APA during the BFU-E to CFU-E transition.

*PABPC1* is a well-known cytoplasmic polyA-binding protein that is expressed in most eukaryotic organisms[36]. Researches have reported that knocking down *PABPC1* can inhibit the proliferation and progression of cancer cells[45–47]. This factor plays a dual role in protecting mRNA polyadenylation and promoting deadenylation, demonstrating a complex regulatory mechanism, and is involved in maintaining transcripts with long 3’UTR isoforms[36, 38]. Previous research has documented that knockdown of *PABPC1* leads to 3’UTR shortening of APA events and inhibits the progression and metastasis of various cancers[36, 47]. In our results, *PABPC1* maintains transcripts stability with long 3’UTR isoforms. Meanwhile, *PABPC1* knockdown results in global APA changes, the inhibition of proliferation and differentiation of early erythroid progenitor cells.

In our downstream analysis, we found *TSC22D1* is one of the important APA genes regulated by *PABPC1*. Further analysis showed that *PABPC1* knockdown led to *TSC22D1* 3’UTR shortening and over-expression. *TSC22D1* can be stimulated by *TGF*β, and regulate the transcription of multiple genes[48]. Inhibition of the *TGF-*β pathway can promote BFU-E expansion[42], thereby suggesting that the altered expression of *TSC22D1* may serve as an important regulator that regulates erythroid progenitor cell expansion. Through further experiments, we discovered that *PABPC1* could influence erythroid progenitor cell expansion, differentiation and survival by regulating APA events and expression of *TSC22D1*(Figure 7).

**Figure 7.**
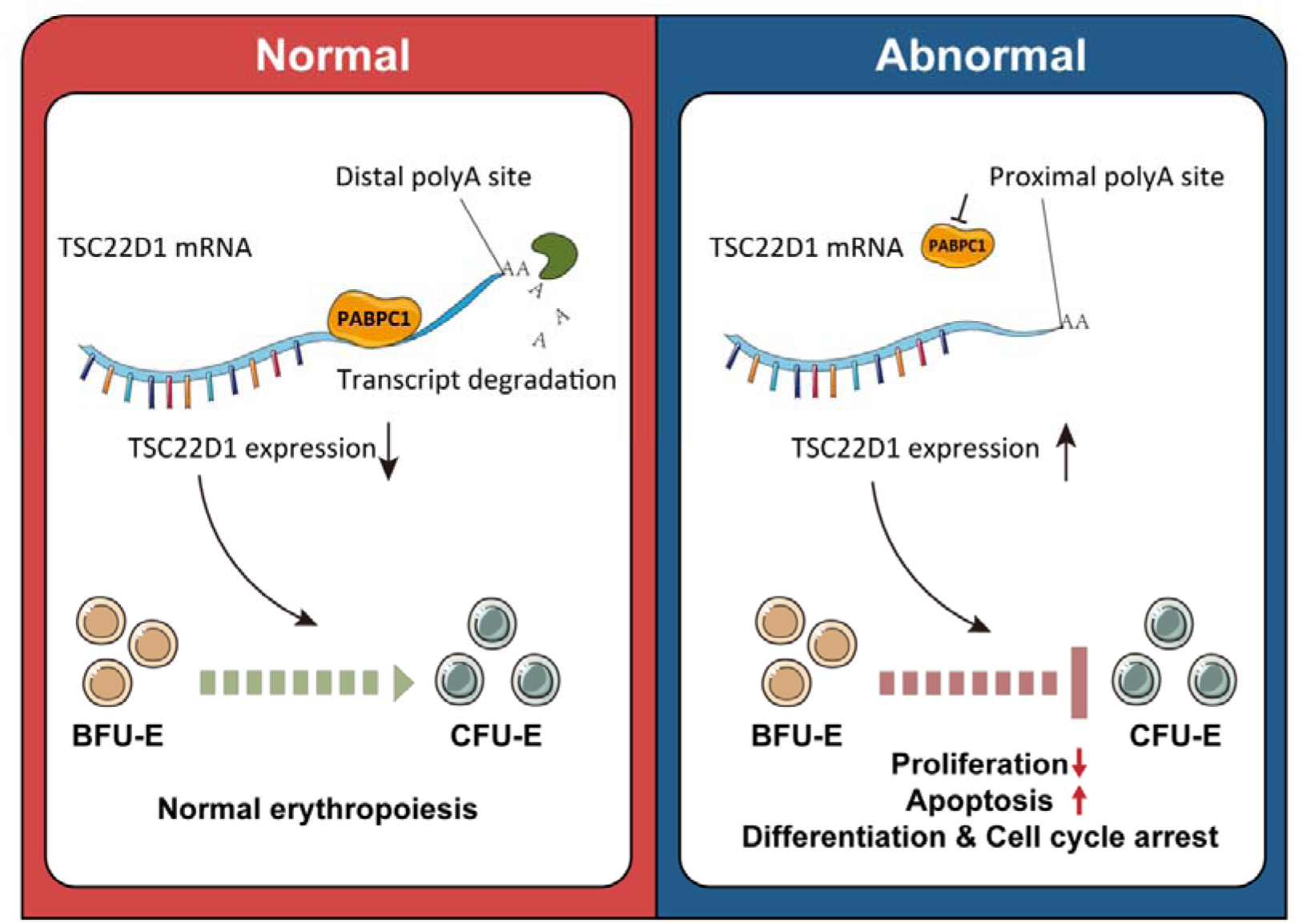
A Model of the Role of Alternative Polyadenylation in early erythropoiesis. PABPC1 control the expression of TSC22D1 by regulating TSC22D1 APA events, thereby inhibiting the expansion and differentiation of early erythroid progenitor cells.

Although the results suggest that APA has a role in erythropoiesis, our study does not directly link APA disruptions to specific erythropoiesis disorders such as PV, MDS and thalassemia. Besides, *PABPC1* is only one of core APA factors involved. Other APA regulatory proteins and their potential interactions with *PABPC1* have not been fully explored. Thus, further studies are essential to map out the complete landscape of APA regulation and its implications for the pathology of erythroid-related diseases.

In conclusion, our study represents a significant advance in the understanding of APA regulation during erythropoiesis, particularly highlighting the pivotal role of *PABPC1*. By demonstrating the impact of APA modulation on the important genes like *TSC22D1*, we provide new insights into the molecular mechanisms governing erythropoiesis. To our knowledge, this is the first study to reveal the substantial impact of APA on erythroid progenitor expansion, enhancing our understanding of erythroid development regulation. We believe these findings could contribute to the broader discourse on erythropoiesis and invite further inquiry into the nuanced role of APA regulation in erythroid disorders.

## Materials and methods

### APA analysis

To explore APA pattern during erythroid differentiation, we collected RNA-Seq data for human early erythropoiesis (BFU-E to CFU-E stages) from GSE61566, and RNA-Seq data for human erythroid terminal differentiation (Pro-E to Ortho-E stages) from GSE53983. Each stage in these datasets contains 3 samples across 7 erythroid stages for a total of 21 samples. We then converted RNA-Seq file from SRA format to fastq format using fastq-dump, and then aligning the sequence to human genome (hg19) by HISAT2. The 21 generated BAM files were then utilized as input for DaPars v2.0, which employs a two-normal mixture model to calculate the percentage of distal polyA site usage index (PDUI), a metric for assessing APA usage for each gene (defined as APA event). PDUI values range from 0 to 1, where higher values indicate increased utilization of distal polyA sites. To ensure the precision of our predictions, we set a coverage threshold for the last exon of ≥30×, following the recommendations of DaPars2. To investigate 3’UTR lengthened/shortened genes during erythropoiesis, we compared PDUI values between different stages, and set the |ΔPDUI| ≥ 0.15 as the criterion. If PDUI at one stage is higher than that of the previous stage, the 3’UTR of the APA event is lengthening during differentiation, and vice versa.

To investigate whether *PABPC1* regulates APA, we analyzed third-generation sequencing data derived from erythroid progenitor cells, including control and *PABPC1* knockout samples. We employed LAPA (https://github.com/mortazavilab/lapa/), a tool specifically designed for third-generation sequencing APA analysis, to systematically assess poly(A) site usage in both the *PABPC1* knockdown (*PABPC1*-KD) and control (NC) samples. Specifically, we selected the two sites exhibiting the greatest variation in usage as the proximal and distal poly(A) sites. Poly(A) site choice was quantified based on the following criteria: (a) Proximal polyA usage choice: An increase in proximal site usage accompanied by a decrease in distal site usage. (b) Distal polyA usage choice: An increase in distal site usage accompanied by a decrease in proximal site usage. Finally, we applied a range of thresholds (±0, ±0.1, ±0.15, and ±0.2) as the criteria to determine poly(A) usage changes.

### Analysis of poly(A) signal motif

A series of cis-elements that regulate alternative polyadenylation are enriched near the poly (A) sites. In order to identify potential cis-elements that affect APA during erythroid development, bedtools was used to extract the genome sequences ±50 bp around proximal and distal polyA sites, respectively. After obtaining the sequences, DREME was used to enrich the features to detect poly(A) signal motifs of proximal and distal polyA sites.

### Prediction of upstream RNA-binding proteins (RBPs) regulating APA

To screen for upstream RBPs regulating APA during erythroid differentiation, we collected a list of reported APA core factors from https://www.nature.com/articles/ncomms6274. The Pearson correlation analysis was employed to assess the relationship between the PDUI values of each APA event and the expression levels of corresponding RBPs. The criteria for determining the significance were stringently defined as FDR ≤ 0.05 and r^2^ ≥ 0.3. Finally, the proportion of APA events linked to RBPs was counted, and RBPs that might affect erythroid differentiation were further screened according to the number of APA events with significant differences.

### Poly(A) Tail Length Analysis

To extract poly(A) tail length information, we employed a script available at https://github.com/ankeetashah/Benchmarking-APA/blob/main/scripts/01_filter_for_p olyA.py, which extracts read IDs, poly(A) tail lengths, and transcript lengths from our ONT files. After extracting the genomic positions of these read IDs from the BAM files, we matched these positions with APA-related poly(A) site peaks identified by LAPA, thereby obtaining poly(A) tail length information for APA genes. Based on our previous LAPA analysis, each APA gene retained two major peaks—defined as distal and proximal sites—and we compared the median poly(A) tail lengths between these sites within the same gene to determine which had longer tails.

### Cell culture

Human mobilized peripheral blood samples were obtained from Xiangya Hospital and the Second Xiangya Hospital of Central South University with approval from the Ethics Committees. Informed consent was obtained from all participating subjects. CD34^+^ cells were purified from human mobilized peripheral blood. The isolated CD34^+^ cells were maintained in a two-stage liquid culture system. During the initial phase (day 0 to day 6), 1×10 /ml CD34^+^ cells were seeded (day 0) in Serum-Free Expansion Medium containing 10% FBS, 1 U/ml EPO, 10 ng/ml IL-3, 50 ng/ml SCF and 0.06 mM α-thioglycerol (Sigma, Catalog No. M6145, Louis, MO). On day 4, fresh medium was added to dilute the cells, and incubation proceeded until day 7. In the subsequent phase (day 7 to day 13), cells were cultured at a density of 1×10 cells/ml in SFEM medium supplemented with 30% FBS, along with 1 U/ml EPO and α-thioglycerol[3].

### Lentivirus infection

PABPC1 shRNAs and control shRNA were obtained from GenePharma Co. Ltd. (Shanghai, China). shRNA targeting sequences: shControl, 5’-TTCTCCGAACGTGTCACGT; shPABPC1 #1, 5’-GGACAAATCCATTGATAAT; shPABPC1 #2, 5’-GAAAGGAGCTCAATGGAAA. shTSC22D1#1, 5’-GAGTTTACCAACTGAGACATT; shTSC22D1#2,

5’-GCAAGCTATGGATCTAGTGAA. Thirty million lentiviruses were used to infect 0.5 million CD34+ cells on day 2. 12 hours after infection, cells were washed with PBS and cultured in new complete medium.

### RNA extraction and qRT-PCR analysis

RNA extracted from primary cultured human erythroid cells using TRIZOL Reagent (Vazyme, Catalog No. R401-01, China) was reverse-transcribed with HiScript II Q RT SuperMix for qPCR (+gDNA wiper) Kit (Vazyme, Catalog No. R223-01, China) according to the manufacturer’s guidelines, and amplified by quantitative polymerase chain reaction (qRT-PCR) with the Cycler (Bio-Rad, Hercules, CA) and appropriate primer pairs. The primers used for qRT-PCR are summarized in Table S3. Relative gene expression was calculated using the 2^-ΔΔCT^ method, with GAPDH used as reference gene. For analysis of APA switches by qRT-PCR, the 2^-ΔCT^ value was calculated. The proximal 2^-ΔCT^ value was divided by the distal 2^-ΔCT^ value, and results were normalized. Results > 1 indicate relative increase of the proximal transcript, whereas results < 1 indicate relative increase of the distal transcript[27]. Primers were designed using NCBI Primer-BLAST.

### Western blotting analysis

For western blotting analysis, whole-cell lysates from cultured cells were prepared with RIPA buffer in the presence of cocktail of protease inhibitor and PhosSTOP phosphatase inhibitor (Roche, Basel, Switzerland). Protein concentration was measured using a Pierce BCA protein assay kit (Thermo Fisher, Catalog No. 23227, Waltham, MA). Protein extracts (30 μg) were boiled, subjected to 10% SDS-PAGE and then transferred to nitrocellulose membranes. The membranes were blocked with 5% nonfat dry milk and incubated with specific primary antibodies (diluted 1:2000) at 4°C overnight. The HRP-conjugated secondary antibodies were used at 1:3000 dilutions for 1.5 h at room temperature. Signals were detected using ECL HRP substrate (Beyotime Biotechnology, China). Western blotting analysis was performed as described previously.

### RNA immunoprecipitation assay

RNA immunoprecipitation (RIP) was performed using a Magna RNA-binding protein immunoprecipitation kit (Millipore, Catalog No.17-700, Bedford, MA) according to the manufacturer’s instructions. Briefly, cells were collected and lysed in complete RIPA buffer containing a protease inhibitor cocktail and RNase inhibitor. Next, the cell lysates were incubated with RIP buffer containing magnetic bead conjugated with human anti-*PABPC1* antibody or control normal rabbit IgG. The samples were digested with proteinase K to isolate the immunoprecipitated RNA. The purified RNA was finally subjected to real-time PCR to demonstrate the presence of the binding targets.

### Plasmids construction, Dual-luciferase reporter assay

We employed the luciferase reporter system to evaluate the impact of *PABPC1* knockdown or overexpression on the regulatory efficiency of functional variants in the proximal or distal regions of the *TSC22D1* gene 3’UTR. The experimental strategy was as follows: First, the proximal and distal regulatory sequences of the *TSC22D1* gene 3’UTR were cloned into the PGL3-BASIC reporter vector and validated by sequencing. Subsequently, functional validation was performed using a dual-luciferase reporter system: 293T cells seeded in 12-well plates were subjected to lentiviral infection to establish stable *TSC22D1* knockdown or overexpression cell models. Following confirmation of gene expression modulation by Western blot, the constructed reporter plasmids and pRL-TK internal control plasmid were co-transfected using Lipofectamine 2000 transfection reagent (Invitrogen, Catalog No.11668019, Carlsbad, CA). Cells were harvested 48 hours post-transfection, and luciferase activity was measured using a dual-luciferase detection system. Quantitative analysis was conducted by calculating the Fluc/Rluc ratio.

### Cell proliferation experiment

After infection with lentivirus, 50000 cells were taken from each group and placed in a 24 well plate for culturing. On days 2, 3, 4, and 5, the cells were gently blown and counted.

### Flow-cytometric analysis

For apoptosis analysis, 5×10^5^ cells were collected, washed by PBS, and resuspended in 100 μl 1×Annexin V Binding buffer (Zeta Life, Catalog No. FL0100, San Francisco, CA). Cells were then stained with 5 μl Annexin V-FITC and 5 μl PI according to the manufacturer’s protocol (Zeta Life, Catalog No. FL0100, San Francisco, CA). After incubating at room temperature for 15 min in the dark, cells were subjected to flow-cytometric analysis on a FACS LSRII flow cytometer (BD Biosciences, Franklin Lakes, NJ). For analysis of early erythroblast differentiation, cells were collected at 48 hours after lentivirus infection. 2×10^5^ cells were washed and resuspended. Cells were then stained with CD45-APC-Cy7, GPA-BV605, CD36-FITC, CD34-APC, CD71-PE, and CD123-PE-Cy7 antibody at 4 for 20 min in the dark. Samples were washed once with PBS and stained with 7AAD-PerCP-Cy5.5 before analysis, then subjected to flow-cytometric analysis on a BD LSR II Flow Cytometer (BD Biosciences, Franklin Lakes, NJ). Unstained cells and APC-Cy7-, BV605-, FITC-, APC-, PE-, PE Cy7-, and PerCP-Cy5.5- were used as negative controls. All flow-cytometric data were analyzed with FlowJo 10.4.

### Colony experiment

Cultured primary CD34+ cells and infected with lentivirus at day 1. were plated in triplicate at a density of 300 cells in 1 ml of MethoCult H4434 classic medium (complete medium containing SCF, IL-3, Epo and GM-CSF) (StemCell, Catalog No.04100, Vancouver, BC) or in 1 ml of MethoCult H4330 medium (StemCell, Catalog No.04100, Vancouver, BC) with Epo only. The CFU-E and BFU-E colonies were defined according to the criteria described by Dover et al. CFU-E colonies were counted on day 7 and BFU-E colonies were counted on day 15.

### Nanopore sequencing

We performed nanopore sequencing (ONT-seq) on early erythroid cells treated with *PABPC1* lentiviral knockdown. The ONT-seq data have been deposited in the NCBI (https://www.ncbi.nlm.nih.gov/) with the dataset identifier PRJNA1061347.

## Authors’ contributions

**Liu H, Gong J, Liang L, Liu J:** Conceptualization, Project administration, Funding acquisition. **Li YN, Yang YB, Liang L:** Writing-original draft. **Li YN, Hu B, Wang Z:** Investigation, Data curation. **Wang W, He XF, Wu XS, Lin S:** Resources. **Yang YB, Li YN:** Formal analysis, Visualization. **Li YN, Yang YB, Liang L, Mohandas N:** Writing-review & editing. All authors have read and approved the final manuscript.

## Acknowledgments

This work was supported by the fundings from National Natural Science Foundation of China (grant numbers 81920108004, 82100137, 81770107, 81702722, 81470362, 81870105, U1804282), National Key Research and Development Program of China (grant number 2108YFA0107800), Hunan Province Clinicalmedical technology innovation guidance project (NO.2021SK50917), Hunan Provincial Clinicalmedical technology innovation guidance project (No. 2023SK4056) and Postgraduate Scientific Research Innovation Project of Hunan Province.

## Supplementary material

**Figure S1 A comprehensive view of 3’UTR length and clustering characteristics of APA events during erythroid differentiation.**

**A**. Boxplot of 3’UTR length variations between transcripts with and without APA events. **B**. 3’UTR size on the x-axis and aUTR size on the y-axis for each APA event, with the correlation coefficient (r) and P value indicated, calculated through Pearson analysis. **C**. 3’UTR size on the x-axis and cUTR size on the y-axis for each APA event, with the correlation coefficient (r) and P value indicated, calculated through Pearson analysis.

**Figure S2 Overexpression of PABPC1 promotes differentiation and expansion of erythroid progenitor cells.**

**A**. Representative flow cytometry chart of early differentiation for early erythroid cell (BFU-E) infected with the same amount of PABPC1 overexpression or control lentivirus. CD34- CD36+ CD71high represent the stage of CFU-E cells, while CD34+ CD36- CD71low represent the stage of BFU-E cells. CD34- CD36+ CD71high and CD34+ CD36- CD71low cells gate from CD45+ GPA- CD123- cells. **B**. Statistical analysis of the CD34- CD36+ CD71high (CFU-E) and CD34+ CD36- CD71low (BFU-E) cells rate(%) from three independent experiments is shown. **C**. Representative images showing the formation of BFU-E and CFU-E colonies (left panel). Statistical analysis of the number of BFU-E and CFU-E colonies formed in colony formation assay (right panel). Data shown are mean ± SD (n = 3) (*P < 0.05, **P < 0.01, ***P < 0.001).

**Figure S3 PABPC4 does not affect the differentiation and expansion of erythroid progenitor cells.**

**A**. RT-qPCR analysis of genes mRNA expression in human BFU-E, CFU-E, and Pro-E populations which sorted by flow cytometry from induced erythroid lineage in vitro. The mRNA expression levels in each group were normalized to GAPDH expression. **B**. RT-qPCR analysis of PABPC4 mRNA expression in early erythroid cell (BFU-E) after infecting with control lentivirus or PABPC4 knockdown lentivirus. GAPDH was used as a loading control. **C**. Early erythroid cell (BFU-E) growth curves determined by cell counting after infecting with control lentivirus or PABPC4 knockdown lentivirus. **D**. Representative flow cytometry chart of early differentiation for early erythroid cell (BFU-E) infected with the same amount of PABPC4 knockdown or control lentivirus. CD34- CD36+ CD71high represent the stage of CFU-E cells, while CD34+ CD36- CD71low represent the stage of BFU-E cells. CD34- CD36+ CD71high and CD34+ CD36- CD71low cells gate from CD45+ GPA- CD123- cells. (E) Statistical analysis of the CD34- CD36+ CD71high (CFU-E) and CD34+ CD36- CD71low (BFU-E) cells rate(%) from three independent experiments is shown. Data shown are mean ± SD (n = 3) (*P < 0.05, **P < 0.01, ***P < 0.001).

**Figure S4 Relationship between alternative polyadenylation and poly(A) tail length.**

**A**. Density distribution of poly(A) tail lengths at proximal (turquoise) and distal (coral) polyadenylation sites in control (NC) and PABPC1 knockdown (PABPC1_KD) samples across three biological replicates. The x-axis represents poly(A) tail length (nucleotides), and the y-axis shows density. **B**. Comparison of poly(A) tail length distributions between control (NC, blue) and PABPC1 knockdown (PABPC1_KD, pink) conditions at distal (top) and proximal (bottom) polyadenylation sites. **C**. Gene count analysis showing APA bias in relation to poly(A) tail length changes. Genes are categorized based on whether they exhibit distal site preference (coral) or proximal site preference (turquoise) across conditions where poly(A) tails were either lengthened or shortened.

**Figure S5 Graphics of ONT coverage tracks comparing Control KD and PABPC1 KD of erythroid progenitor cells.**

**A**. TSC22D1, **B**. PTP4A2, **C**. P2RX1, and **D**. PARP1. E. The heatmap represents the mean relative expression of TSC22D1 mRNA isoform utilizing proximal and distal polyA sites based on qRT-PCR analysis (PABPC1 knockdown versus control group).

**Figure S6 Western blotting of PABPC1 expression upon knockdown or overexpression in human induced BFU-E.**

**A**. Representative images of western blotting showing PABPC1 expression after the PABPC1 knockdown in human induced BFU-E in vitro. **B**. Representative images of western blotting showing PABPC1 expression after the PABPC1 overexpression in human induced BFU-E in vitro.

**Figure S7 TSC22D1 mRNA expression from GSE61566 and GSE53983.**

**A**. The mRNA expression of TSC22D1 from RNA-seq data GSE61566 and GSE53983.

**Table S1 APA Changes in PABPC1 Knockdown Across Different Thresholds**

**Table S2 Comparison of Poly(A) Tail Lengths Between Proximal and Distal APA Sites**

**Table S3 The primers used for qRT-PCR**

**Table S4 The median polyA tail lengths for gene distal and proximal APA sites across ONT samples**

## References

[1] Yamamoto R, Morita Y, Ooehara J, Hamanaka S, Onodera M, Rudolph KL, et al. Clonal analysis unveils self-renewing lineage-restricted progenitors generated directly from hematopoietic stem cells. Cell 2013;154:1112–26.

[2] Dong F, Hao S, Zhang S, Zhu C, Cheng H, Yang Z, et al. Differentiation of transplanted haematopoietic stem cells tracked by single-cell transcriptomic analysis. Nat Cell Biol 2020;22:630–9.

[3] Li J, Hale J, Bhagia P, Xue F, Chen L, Jaffray J, et al. Isolation and transcriptome analyses of human erythroid progenitors: BFU-E and CFU-E. Blood 2014;124:3636–45.

[4] Li Y, Zhang H, Hu B, Wang P, Wang W, Liu J. Post-transcriptional regulation of erythropoiesis. Blood Sci 2023;5:150–9.

[5] Han X, Zhang J, Peng Y, Peng M, Chen X, Chen H, et al. Unexpected role for p19INK4d in posttranscriptional regulation of GATA1 and modulation of human terminal erythropoiesis. Blood 2017;129:226–37.

[6] Peng Y, Tang L, Li Y, Song J, Liu H, Wang P, et al. Comprehensive proteomic analysis reveals dynamic phospho-profiling in human early erythropoiesis. Br J Haematol 2022;199:427–42.

[7] Iskander D, Psaila B, Gerrard G, Chaidos A, En Foong H, Harrington Y, et al. Elucidation of the EP defect in Diamond-Blackfan anemia by characterization and prospective isolation of human EPs. Blood 2015;125:2553–7.

[8] Nandakumar SK, Ulirsch JC, Sankaran VG. Advances in understanding erythropoiesis: evolving perspectives. Br J Haematol 2016;173:206–18.

[9] Hellstrom-Lindberg E, Tobiasson M, Greenberg P. Myelodysplastic syndromes: moving towards personalized management. Haematologica 2020;105:1765–79.

[10] Cazzola M. Ineffective erythropoiesis and its treatment. Blood 2022;139:2460–70.

[11] Gao X, Lee HY, da Rocha EL, Zhang C, Lu YF, Li D, et al. TGF-beta inhibitors stimulate red blood cell production by enhancing self-renewal of BFU-E erythroid progenitors. Blood 2016;128:2637–41.

[12] Zhang L, Prak L, Rayon-Estrada V, Thiru P, Flygare J, Lim B, et al. ZFP36L2 is required for self-renewal of early burst-forming unit erythroid progenitors. Nature 2013;499:92–6.

[13] Gruber AJ, Zavolan M. Alternative cleavage and polyadenylation in health and disease. Nat Rev Genet 2019;20:599–614.

[14] Mitschka S, Mayr C. Context-specific regulation and function of mRNA alternative polyadenylation. Nat Rev Mol Cell Biol 2022;23:779–96.

[15] Tian B, Manley JL. Alternative polyadenylation of mRNA precursors. Nat Rev Mol Cell Biol 2017;18:18–30.

[16] Zhang Y, Liu L, Qiu Q, Zhou Q, Ding J, Lu Y, et al. Alternative polyadenylation: methods, mechanism, function, and role in cancer. J Exp Clin Cancer Res 2021;40:51.

[17] Brumbaugh J, Di Stefano B, Wang X, Borkent M, Forouzmand E, Clowers KJ, et al. Nudt21 Controls Cell Fate by Connecting Alternative Polyadenylation to Chromatin Signaling. Cell 2018;172:106–20 e21.

[18] Qin H, Ni H, Liu Y, Yuan Y, Xi T, Li X, et al. RNA-binding proteins in tumor progression. J Hematol Oncol 2020;13:90.

[19] Blake D, Lynch KW. The three as: Alternative splicing, alternative polyadenylation and their impact on apoptosis in immune function. Immunol Rev 2021;304:30–50.

[20] Witkowski MT, Lee S, Wang E, Lee AK, Talbot A, Ma C, et al. NUDT21 limits CD19 levels through alternative mRNA polyadenylation in B cell acute lymphoblastic leukemia. Nat Immunol 2022;23:1424–32.

[21] Xia Z, Donehower LA, Cooper TA, Neilson JR, Wheeler DA, Wagner EJ, et al. Dynamic analyses of alternative polyadenylation from RNA-seq reveal a 3’-UTR landscape across seven tumour types. Nat Commun 2014;5:5274.

[22] Lee SH, Singh I, Tisdale S, Abdel-Wahab O, Leslie CS, Mayr C. Widespread intronic polyadenylation inactivates tumour suppressor genes in leukaemia. Nature 2018;561:127–31.

[23] Xiang Y, Ye Y, Lou Y, Yang Y, Cai C, Zhang Z, et al. Comprehensive Characterization of Alternative Polyadenylation in Human Cancer. J Natl Cancer Inst 2018;110:379–89.

[24] Wang L, Lang GT, Xue MZ, Yang L, Chen L, Yao L, et al. Dissecting the heterogeneity of the alternative polyadenylation profiles in triple-negative breast cancers. Theranostics 2020;10:10531–47.

[25] Miles WO, Lembo A, Volorio A, Brachtel E, Tian B, Sgroi D, et al. Alternative Polyadenylation in Triple-Negative Breast Tumors Allows NRAS and c-JUN to Bypass PUMILIO Posttranscriptional Regulation. Cancer Res 2016;76:7231–41.

[26] Song J, Nabeel-Shah S, Pu S, Lee H, Braunschweig U, Ni Z, et al. Regulation of alternative polyadenylation by the C2H2-zinc-finger protein Sp1. Mol Cell 2022;82:3135–50 e9.

[27] Sommerkamp P, Altamura S, Renders S, Narr A, Ladel L, Zeisberger P, et al. Differential Alternative Polyadenylation Landscapes Mediate Hematopoietic Stem Cell Activation and Regulate Glutamine Metabolism. Cell Stem Cell 2020;26:722–38 e7.

[28] Kini HK, Kong J, Liebhaber SA. Cytoplasmic poly(A) binding protein C4 serves a critical role in erythroid differentiation. Mol Cell Biol 2014;34:1300–9.

[29] van Zalen S, Lombardi AA, Jeschke GR, Hexner EO, Russell JE. AUF-1 and YB-1 independently regulate beta-globin mRNA in developing erythroid cells through interactions with poly(A)-binding protein. Mech Dev 2015;136:40–52.

[30] Feng X, Li L, Wagner EJ, Li W. TC3A: The Cancer 3’ UTR Atlas. Nucleic Acids Res 2018;46:D1027–D30.

[31] Ielasi FS, Ternifi S, Fontaine E, Iuso D, Coute Y, Palencia A. Human histone pre-mRNA assembles histone or canonical mRNA-processing complexes by overlapping 3’-end sequence elements. Nucleic Acids Res 2022;50:12425–43.

[32] Liu X, Hoque M, Larochelle M, Lemay JF, Yurko N, Manley JL, et al. Comparative analysis of alternative polyadenylation in S. cerevisiae and S. pombe. Genome Res 2017;27:1685–95.

[33] Beaudoing E, Freier S, Wyatt JR, Claverie JM, Gautheret D. Patterns of variant polyadenylation signal usage in human genes. Genome Res 2000;10:1001–10.

[34] Kwon B, Fansler MM, Patel ND, Lee J, Ma W, Mayr C. Enhancers regulate 3’ end processing activity to control expression of alternative 3’UTR isoforms. Nat Commun 2022;13:2709.

[35] Li W, You B, Hoque M, Zheng D, Luo W, Ji Z, et al. Systematic profiling of poly(A)+ transcripts modulated by core 3’ end processing and splicing factors reveals regulatory rules of alternative cleavage and polyadenylation. PLoS Genet 2015;11:e1005166.

[36] Zhang Y, Chen C, Liu Z, Guo H, Lu W, Hu W, et al. PABPC1-induced stabilization of IFI27 mRNA promotes angiogenesis and malignant progression in esophageal squamous cell carcinoma through exosomal miRNA-21-5p. J Exp Clin Cancer Res 2022;41:111.

[37] Gray NK, Hrabalkova L, Scanlon JP, Smith RW. Poly(A)-binding proteins and mRNA localization: who rules the roost? Biochem Soc Trans 2015;43:1277–84.

[38] Kuhn U, Gundel M, Knoth A, Kerwitz Y, Rudel S, Wahle E. Poly(A) tail length is controlled by the nuclear poly(A)-binding protein regulating the interaction between poly(A) polymerase and the cleavage and polyadenylation specificity factor. J Biol Chem 2009;284:22803–14.

[39] Apponi LH, Leung SW, Williams KR, Valentini SR, Corbett AH, Pavlath GK. Loss of nuclear poly(A)-binding protein 1 causes defects in myogenesis and mRNA biogenesis. Hum Mol Genet 2010;19:1058–65.

[40] Deamer D, Akeson M, Branton D. Three decades of nanopore sequencing. Nat Biotechnol 2016;34:518–24.

[41] Wang Y, Zhao Y, Bollas A, Wang Y, Au KF. Nanopore sequencing technology, bioinformatics and applications. Nat Biotechnol 2021;39:1348–65.

[42] Dragotto J, Canterini S, Del Porto P, Bevilacqua A, Fiorenza MT. The interplay between TGF-beta-stimulated TSC22 domain family proteins regulates cell-cycle dynamics in medulloblastoma cells. J Cell Physiol 2019;234:18349–60.

[43] Canterini S, Carletti V, Nusca S, Mangia F, Fiorenza MT. Multiple TSC22D4 iso-/phospho-glycoforms display idiosyncratic subcellular localizations and interacting protein partners. FEBS J 2013;280:1320–9.

[44] Zheng Z, Chen X, Cai X, Lin H, Xu J, Cheng X. RNA-binding protein MEX3D promotes cervical carcinoma tumorigenesis by destabilizing TSC22D1 mRNA. Cell Death Discov 2022;8:250.

[45] Li J, Pei M, Xiao W, Liu X, Hong L, Yu Z, et al. The HOXD9-mediated PAXIP1-AS1 regulates gastric cancer progression through PABPC1/PAK1 modulation. Cell Death Dis 2023;14:341.

[46] YuFeng Z, Ming Q. Expression and prognostic roles of PABPC1 in hepatocellular carcinoma. Int J Surg 2020;84:3–12.

[47] Qi Y, Wang M, Jiang Q. PABPC1--mRNA stability, protein translation and tumorigenesis. Front Oncol 2022;12:1025291.

[48] Lu Y, Kitaura J, Oki T, Komeno Y, Ozaki K, Kiyono M, et al. Identification of TSC-22 as a potential tumor suppressor that is upregulated by Flt3-D835V but not Flt3-ITD. Leukemia 2007;21:2246–57.

