## Supplemental Table 1 for "Regulation of Alternative Polyadenylation Events by PABPC1 Affects Erythroid Progenitor Cell Expansion"

Table S1. APA Changes in PABPC1 Knockdown Across Different Thresholds

| **Threshold of polyA Site Usage (PABPC1-KD/NC)** | **Distal Usage Choice** | **Proximal Usage Choice** |
| --- | --- | --- |
| ± 0 | 3561 | 4049 |
| ± 0.1 | 1761 | 2000 |
| ± 0.15 | 1132 | 1268 |
| ± 0.2 | 766 | 846 |
