## Supplemental Table 2 for "Regulation of Alternative Polyadenylation Events by PABPC1 Affects Erythroid Progenitor Cell Expansion"

Table S2. Comparison of Poly(A) Tail Lengths Between Proximal and Distal APA Sites

| **Sample ID** | **Proximal site with Longer polyA Tails** | **Distal site with Longer polyA Tails** |
| --- | --- | --- |
| NC_1 | 2,968 | 3,417 |
| NC_2 | 3,036 | 3,390 |
| NC_3 | 2,854 | 3,475 |
| PABPC1_KD_1 | 2,710 | 3,298 |
| PABPC1_KD_2 | 3,090 | 3,584 |
| PABPC1_KD_3 | 2,923 | 3,551 |
