## Supplemental Table 3 for "Regulation of Alternative Polyadenylation Events by PABPC1 Affects Erythroid Progenitor Cell Expansion"

| Table S3. The primers used for qRT-PCR | |
| --- | --- |
| Target gene | Primer sequence (5'→ 3') |
| GAPDH-F | GGAGCGAGATCCCTCCAAAAT |
| GAPDH-R | GGCTGTTGTCATACTTCTCATGG |
| PABPC1-F | AGCAAATGTTGGGTGAACGG |
| PABPC1-R | ACCGGTGGCACTGTTAACTG |
| F-PARP1-P | GCCGAGATCATCAGGAAGTATG |
| R-PARP1-P | ATTCGCCTTCACGCTCTATC |
| F-PARP1-D | GAGCTTTCCTTCTCCAGGAATA |
| R-PARP1-D | AGCCCTTGGGTAAGTATATTTGT |
| F-P2RX1-P | ACCATCGGCTCTGGAATTG |
| R-P2RX1-P | CTGCTTGTAGTAGTGCCTCTTAG |
| F-P2RX1-D | GCTCAGTAGATACGTGTGGTTAG |
| R-P2RX1-D | AGGAAGAAACAGGGCTCAAG |
| F-TSC22D1-P | ATTTGATGTATGCGGTCAGAGA |
| R-TSC22D1-P | AGCAGATTGTTCTCCTGCTC |
| F-TSC22D1-D | TAGAGCTGGTGGGAGATTGA |
| R-TSC22D1-D | GCCCGTGTCACCACTTTAT |
| F-DDX17-P | CTGATAGGCAGACACTGATGTG |
| R-DDX17-P | CCTACGTTGATCTGGGTGTAATC |
| F-DDX17-D | CGTTGTGTGGACTTCTCATCTAA |
| R-DDX17-D | ATCCTGTGCCTGAACATCAC |
| F-RSL1D1-P | CTCCCTCCAAACTCTAAAGAAGG |
| R-RSL1D1-P | CGCCTAATTCTGGCATCAGTA |
| F-RSL1D1-D | GTCTGACTTGAGGTCTGTGATG |
| R-RSL1D1-D | CCACTCTGGGAATGACTACAAA |
| F-PTP4A2-P | TGTTGCAGGATTGGGAAGAA |
| R-PTP4A2-P | ATGCCCATTGGTATCTCTGAAG |
| F-PTP4A2-D | AGCCAGTTTGACATAGAGAGATG |
| R-PTP4A2-D | AGGGTTCAGGGTTTGGTTT |
| F-PSMA5-P | CCCAAGCTGTGTCCAATCT |
| R-PSMA5-P | GGTCCTTTCTCATCAACTCCTC |
| F-PSMA5-D | GCACCTACCTACTATATGCCAAG |
| R-PSMA5-D | CCCACTAGAAGTTTCACAGGAG |
| F-ZFP36L2-P | CACTTCTGTCCGCCTTCTAC |
| R-ZFP36L2-P | CATGTTGTTCAGGTTGAGGTTG |
| F-ZFP36L2-D | TCCATGTAGAAAGGCAGGAATG |
| R-ZFP36L2-D | TCTGTCCAAAGTAGTCCAGAGA |
| F-PAPOLA | GAGGGATGCCAGATGTTGATAA |
| R-PAPOLA | GGAGCACAATGGAGGATTCATA |
| F-PABPN1 | GAGCTACAGAACGAGGTAGAGA |
| R-PABPN1 | CATAGATGGAACGGGCATCAG |
| F-PAPOLG | TGCCATTGGAGGAGAATCTATG |
| R-PAPOLG | GCTGGAACTGGAGATGATGAA |
| F-PPP1CA | GAGACGCTACAACATCAAACTG |
| R-PPP1CA | AGATCTTTTCGTCCACTATGGC |
| F-CPSF2 | TCTGCCCTTTGCTATCTTCTC |
| R-CPSF2 | TGCTTCCTCAGGGAATCAATAA |
| F-CSTF2T | GTGTAATGGAGAGGAGAGGAATG |
| R-CSTF2T | CCCTGAATTCCACCAGTCATAG |
| F-FIP1L1 | CAGCAAGCAGTGGGACTATTA |
| R-FIP1L1 | TCTCTGGTGCGTTCTCTTTC |
| F-WDR33 | GACCACGGAGGATATGTGAAAT |
| R-WDR33 | TTGTGTATAAACCTGGCCTCTC |
| F-CPSF7 | TCCCAAGCCCAACAACAA |
| R-CPSF7 | GGAGAAGCTGCCCACATAAA |
| F-CPSF6 | CCCTGGAAGGGAAATGGATAC |
| R-CPSF6 | ATCAGACACAGCTCTCGAAATAG |
| F-SYMPK | AGCAAGCTCACATCCCTAAC |
| R-SYMPK | TCCAGATCATCCTCCTCCAA |
| F-PCF11 | CCCTGGATGTCAGAGTCAATTC |
| R-PCF11 | GCTCCTCGGGCGATTTATTTA |
| F-CPSF3 | ACTAGCCAAGGTTATGGGATTT |
| R-CPSF3 | AGGTCGCAAGGAGAAAGTATG |
| F-CSTF1 | ATAAAGGACCATGCCGTGTAG |
| R-CSTF1 | AACATCCTCTCTGTGTCAAGTATC |
| F-PPP1CB | ATCTTTCTCAGCCAGCCTATTC |
| R-PPP1CB | CTGGTGGGAAACCTCCATATTC |
| F-NUDT21 | TGGAATGAGGAGGACTGTAGAA |
| R-NUDT21 | CACCACCAGGTAGTTTGAAGAA |
| F-CSTF2 | CTGAGGTTGGACCTGTTGTTAG |
| R-CSTF2 | GCTGTCTCTTGGTCTTGGTATT |
| F-CPSF4 | CATCAGTGGTGAGAAGACAGTT |
| R-CPSF4 | GGGCATCTTGGTCATGTCATA |
| F-CPSF1 | TGAGACCATCGAGAGAGATGAG |
| R-CPSF1 | CGATCCTGGCATTGGGAATAG |
| F-CSTF3 | GTCCCAGAGAAGGTGAAGAAAG |
| R-CSTF3 | TGATTCTGTGCCTCTCGAATG |
| F-PABPC4 | CTCCAGCTCCTGCCAATTTA |
| R-PABPC4 | TAGTCGTCGCTGCTCTCTAT |
| F-RBBP6 | CTCCAGCTCCTGCCAATTTA |
| R-RBBP6 | TAGTCGTCGCTGCTCTCTAT |
